# hypeR: An R Package for Geneset Enrichment Workflows

**DOI:** 10.1101/656637

**Authors:** Anthony Federico, Stefano Monti

## Abstract

**Summary:** Geneset enrichment is a popular method for annotating high-throughput sequencing data. Existing tools fall short in providing the flexibility to tackle the varied challenges researchers face in such analyses, particularly when analyzing many signatures across multiple experiments. We present a comprehensive R package for geneset enrichment workflows that offers multiple enrichment, visualization, and sharing methods in addition to novel features such as hierarchical geneset analysis and built-in markdown reporting. hypeR is a one-stop solution to performing geneset enrichment for a wide audience and range of use cases.

**Availability and implementation:** The most recent version of the package is available at https://github.com/montilab/hypeR.

**Supplementary information:** Comprehensive documentation and tutorials, are available at https://montilab.github.io/hypeR-docs.

## 1 INTRODUCTION

Geneset enrichment is an important step in biological data analysis workflows, particularly in bioinformatics and computational biology. At a basic level, one is performing a hypergeometric or Kolmogorov–Smirnov test to determine if a group of genes is over-represented or enriched, respectively, in pre-defined sets of genes, which suggests some biological relevance. The R package hypeR brings a fresh take to geneset enrichment, focusing on the analysis, visualization, and reporting of enriched genesets. While similar tools exists - such as Enrichr (Kuleshov et al., 2016), fgsea (Sergushichev, 2016), and clusterProfiler (Wang et al., 2012), among others - hypeR excels in the downstream analysis of geneset enrichment workflows – in addition to sometimes overlooked upstream analysis methods such as allowing for a flexible background population size or reducing genesets to a background distribution of genes. Finding relevant biological meaning from a large number of often obscurely labeled genesets may be challenging for researchers. hypeR overcomes this barrier by incorporating hierarchical ontologies - also referred to as relational genesets - into its workflows, allowing researchers to visualize and summarize their data at varying levels of biological resolution. All analysis methods are compatible with hypeR’s markdown features, enabling concise and reproducible reports easily shareable with collaborators. Additionally, users can import custom genesets that are easily defined, extending the analysis of genes to other areas of interest such as proteins, microbes, metabolites, etc. The hypeR package goes beyond performing basic enrichment, by providing a suite of methods designed to make routine geneset enrichment seamless for scientists working in R.

## 2 IMPLEMENTATION

hypeR is a Bioconductor package written completely in R. The core function hypeR() accepts one or more signatures and a list of genesets to test for either over-representation or enrichment, depending on whether the signature is a vector of unranked genes, or a ranked vector of genes with or without weights. The former case is applicable to clusters of genes, such as those identified through co-expression analysis, while the latter is useful when signatures of genes can be ranked, such as through differential expression analysis. Despite its flexibility, hypeR() always returns one or more hyp objects that are defined using R6 (Mailund & Mailund, 2017), which is an implementation of encapsulated object-oriented programming for R. A hyp object contains all information relevant to the enrichment analysis, including a data frame of results, enrichment plots for each geneset tested, as well as the arguments used to perform the analysis. All downstream functions used for analysis, visualization, and reporting recognize hyp objects and utilize their data. Adopting an object oriented framework brings modularity to hypeR, enabling flexible workflows. Additionally, most of hypeR’s functionalities are applicable after enrichment results have been calculated. Therefore, users can perform enrichment with other popular tools, and use hypeR to analyze the results, by formatting the output into a hyp object. As an example, the documentation includes a tutorial for transforming the output of fgsea into a hyp object and analyzing the data with hypeR.

## 3 APPLICATION

hypeR() requires two arguments, a signature of genes and a list of genesets. Depending on the type of signature and genesets provided, downstream functions will behave differently. The simplest use case involves a single signature. When running hypeR() on a single signature, one hyp object is returned. One can extract the data slot to manually inspect the results, or use hyp-compatible functions in their analysis.

### 3.1 Visualization Plots

To visualize hyp objects, users can call hyp_show() to view the data as an interactive table, hyp_dots() to plot top enriched genesets, and hyp_emap() to represent them as an enrichment map. hyp_dots() returns a horizontal dot plot whereby each dot represents a geneset, colored by its enrichment significance and scaled by its size. hyp_emap() returns an interactive enrichment network generated with visNetwork. An enrichment map is useful for identifying clusters of related genesets. Each node represents a geneset, colored by its enrichment significance, and each edge represents the Jaccard index or overlap similarity of two genesets.

### 3.2 Relational Genesets

When dealing with hundreds of genesets, it is useful to summarize results into more general and interpretable biological themes. To aid this process, hypeR() recognizes a special type of genesets called relational genesets (rgsets). An rgsets object is an R6 class which organizes genesets by curated relationships. For example, Reactome (Fabregat et al., 2018) defines their genesets as a hierarchy of pathways, which can be coerced into an rgsets object. Relational genesets have three data attributes including gsets, nodes, and edges. The gsets attribute includes the geneset information for the leaf nodes of the hierarchy, the nodes attribute describes all nodes in the hierarchy, including internal nodes, and the edges attribute describes the hierarchy edges. One can visualize the enriched genesets in a hierarchy map with hyp_hmap(). Each node represents a geneset, where the shade of the gold border indicates the enrichment significance while each edge represents a directed relationship between genesets in the hierarchy (Fig. 1B). Users can double click internal nodes to cluster their first degree connections, allowing for dynamic rescaling of the network to higher or lower degrees of complexity.

**Fig. 1.**
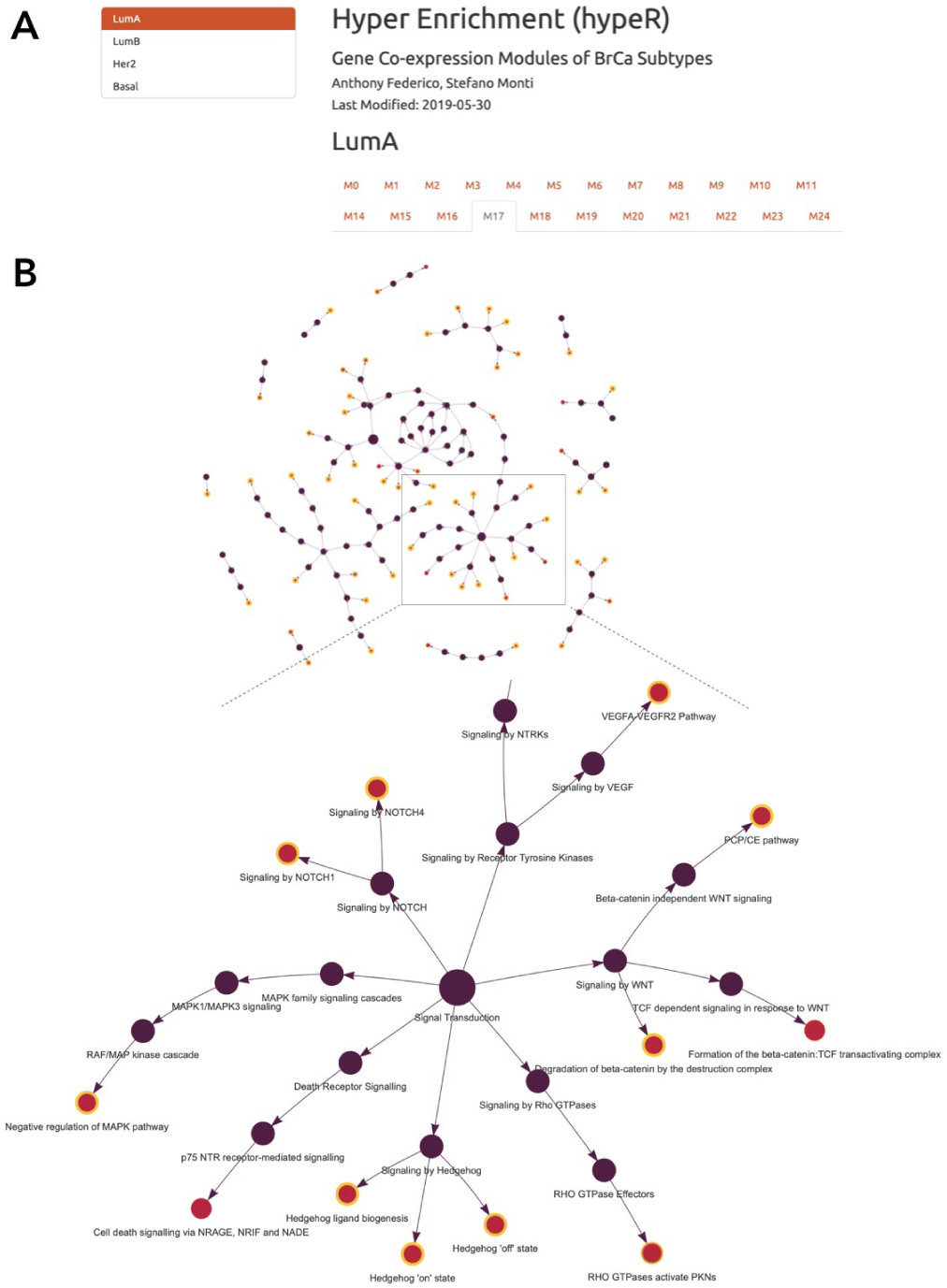
(**A**) An example of markdown reporting for multiple signatures across multiple experiments. (**B**) A hierarchy of enriched genesets visualized with hypeR.

### 3.3 Reporting

After a hypeR session, hyp objects can be saved as R data files, allowing users to easily resume their analysis later. Additionally, hyp objects can be exported to Excel or other tabular formats. Lastly, hypeR allows for generating comprehensive markdown reports. This feature is most useful in cases where there may be multiple signatures across multiple experiments (Fig. 1A). In this case, each experiment is a section of the report, and each signature within that experiment is displayed under a tab. Within each tab can be a combination of interactive tables, summary plots, enrichment maps, and hierarchy maps.

The ability to process, interpret, and share large amounts of biological data is exemplified in Figure 1, where gene co-expression modules were computed from samples of four breast cancer subtypes (LumA, LumB, Her2, and Basal) with RNA-seq data from TCGA-BRCA, resulting in hundreds of signatures to annotate. The markdown report allows researchers to efficiently browse signatures and to annotate them using a hierarchy of enriched genesets. For example, Module 17 of LumA samples is enriched by three genesets related to the Hedgehog signaling pathway, namely, Hedgehog Ligand Biogenesis, Hedgehog *on* State, and Hedgehog *off* State. If a more general theme is desired, the user can use the parent node of these genesets, annotating the enrichment simply as Signaling by Hedgehog. Overall, this style of geneset enrichment workflows greatly reduces the manual labor and interpretation by researchers.

### 3.4 Data

By default, hypeR provides a method for loading MSigDB (Liberzon et al., 2011) genesets via the R package msigdb. hypeR also enables users to download hundreds of other popular genesets that are hosted in a hypeR-compatible format by <u>hypeR-db on GitHub</u>. Again, because hypeR accepts genesets in the form of a named list of character vectors, users can provide any genesets desired.

## 4 CONCLUSION

hypeR is an open source R package, released under the GNU license, available for Linux, Mac OS, and Windows on Bioconductor or GitHub. Comprehensive documentation and tutorials are available at https://montilab.github.io/hypeR-docs. In summary, we provide an easy-to-use tool to improve the efficiency of performing geneset enrichment workflows for a variety of research needs.

## ACKNOWLEDGEMENTS

We would like to thank dbGap (phs000178.v9.p8) for granting access to the TCGA data used to generate Fig. 1.

## REFERENCES

Fabregat, A., Jupe, S., Matthews, L., Sidiropoulos, K., Gillespie, M., Garapati, P., … D’Eustachio, P. (2018). The Reactome Pathway Knowledgebase. Nucleic Acids Research. https://doi.org/10.1093/nar/gkx1132

Kuleshov, M. V, Jones, M. R., Rouillard, A. D., Fernandez, N. F., Duan, Q., Wang, Z., … Ma’ayan, A. (2016). Enrichr: a comprehensive gene set enrichment analysis web server 2016 update. Nucleic Acids Research. https://doi.org/10.1093/nar/gkw377

Liberzon, A., Subramanian, A., Pinchback, R., Thorvaldsdóttir, H., Tamayo, P., & Mesirov, J. P. (2011). Molecular signatures database (MSigDB) 3.0. Bioinformatics. https://doi.org/10.1093/bioinformatics/btr260

Mailund, T., & Mailund, T. (2017). R6 Classes. In Advanced Object-Oriented Programming in R. https://doi.org/10.1007/978-1-4842-2919-4_7

Sergushichev, A. A. (2016). An algorithm for fast preranked gene set enrichment analysis using cumulative statistic calculation. BioRxiv. https://doi.org/10.1101/060012

Yu, G., Wang, L.-G., Han, Y., & He, Q.-Y. (2012). clusterProfiler: an R Package for Comparing Biological Themes Among Gene Clusters. OMICS: A Journal of Integrative Biology. https://doi.org/10.1089/omi.2011.0118

